# The Immune Deficiency Pathway Regulates Metabolic Homeostasis in *Drosophila*

**DOI:** 10.1101/253542

**Authors:** Saeideh Davoodi, Anthony Galenza, Andrew Panteluk, Rujuta Deshpande, Meghan Ferguson, Savraj Grewal, Edan Foley

## Abstract

Immune and metabolic pathways collectively influence host responses to microbial invaders, and mutations in one pathway frequently disrupt activity in the other. We used the *Drosophila* model to characterize metabolic homeostasis in flies with modified Immune Deficiency (IMD) pathway activity. The IMD pathway is very similar to the mammalian Tumor Necrosis Factor-alpha pathway, a key regulator of vertebrate immunity and metabolism. We found that persistent activation of IMD resulted in hyperglycemia, depleted fat reserves, and developmental delays, implicating IMD in metabolic regulation. Consistent with this hypothesis, we found that *imd* mutants weigh more, are hyperlipidemic, and have impaired glucose tolerance. To test the importance of metabolic regulation for host responses to bacterial infection, we challenged insulin pathway mutants with lethal doses of several *Drosophila* pathogens. We found that loss-of-function mutations in the insulin pathway impacted host responses to infection in a manner that depends on the route of infection, and the identity of the infectious microbe. Combined, our results support a role for coordinated regulation of immune and metabolic pathways in host containment of microbial invaders.

## INTRODUCTION

The gastrointestinal tract processes ingested material in a manner that prevents microbial penetration of the host interior, and allows an orderly flow of essential nutrients to metabolic organs. Once inside the host, nutrients initiate signal transduction pathways that control growth and development. Prominent metabolic regulators appeared very early during animal evolution - an estimated billion years ago in the case of the insulin peptides (1), and execute conserved functions across the animal kingdom. For example, insulin peptides control the uptake and storage of nutrients, activate cellular growth, and influence longevity in animals as diverse as worms, flies, and rodents (2–4).

Metabolism undergoes a fundamental shift upon infection (5). At this point, germline-encoded pattern recognition receptors detect the molecular signatures of alien microbes, and initiate physiological responses designed to neutralize the invader, and optimize host survival. We tend to focus on immunity as the generation of molecules that eliminate the invader and eradicate infected cells. However, microbial detection initiates a complex spectrum of responses, that may include elements as diverse as increased body temperature, lethargy, loss of appetite, social isolation, and tolerance mechanisms that neutralize pathogens without affecting their numbers (6). Metabolic adaptations are a common theme in the host response to infection (7). In this case, hosts balance traditional metabolic needs against the immediate threat presented by the microbe, and alter metabolic pathway activity accordingly.

The integration of immune and metabolic pathways is particularly apparent in insects such as *Drosophila melanogaster*, where the fat body simultaneously regulates energy storage and humoral immunity. Under optimal conditions, the larval fat body detects circulating sugars and amino acids in the hemolymph to control the release of *Drosophila* Insulin-Like Peptides (dILPs) from the brain (8). dILPs enter circulation and orchestrate the actions of metabolic organs such as muscle, and fat. At the same time, pattern recognition receptors survey the hemolymph for microbe-associated molecular patterns (MAMPs) that indicate infection. Several types of MAMP activate Toll-mediated responses in the fat body (9), while the Immune Deficiency (IMD) pathway, an evolutionary relative of the Tumor Necrosis Factor (TNF) pathway (10), responds to bacterial diaminopimelic-acid containing peptidoglycan (11). The host integrates signals from immune and metabolic pathways to determine the net output of the fat body. For example, under times of high nutrient availability and limited microbial detection, the *Drosophila* MEF2 transcription factor is phosphorylated, and promotes lipogenesis and glycogenesis, molecular pathways that support growth in the animal (12). However, bacterial infection causes a loss of MEF2 phosphorylation, an event that shifts fat body activity from the accumulation of energy stores to the release of antimicrobial peptides. Such metabolic shifts, are common in *Drosophila* responses to infection, and frequently include alterations to the activity of insulin/target of rapamycin (TOR) pathway elements (13, 14).

Molecular links between immune and metabolic pathways are conserved across vast evolutionary distances, and abnormal immune-metabolic signals are linked to several pathological states. For example, inflammation is involved in the development of chronic metabolic disorders such as insulin resistance, and type two diabetes (15, 16). In experimental models of obesity, adipose tissue-resident macrophages produce TNF (17, 18), and TNF contributes to the development of obesity-induced insulin resistance (19–21). Indeed, treatment with anti-inflammatory salicylates improves obesity-induced insulin resistance, and type two diabetes (22, 23). However, despite the impact of inflammatory cues on metabolic homeostasis, we do not fully understand how the respective pathways communicate.

We used the *Drosophila* model to characterize the contributions of IMD to immune-metabolic homeostasis. We found that activation of IMD in the fat body has the molecular, genetic, and phenotypic signatures of alterations to host metabolism. Transcriptionally, activation of IMD resulted in a gene expression signature consistent with diminished insulin/TOR activity. Physiologically, IMD activation caused a depletion of lipid stores, hyperglycemia, delayed development, and a reduction in adult size. In follow-up studies, we found that loss of function *imd* mutants weigh more, have deregulated insulin signaling, hyperlipidemia, and impaired glucose tolerance. The apparent links between IMD and metabolism led us to speculate that loss of key of metabolic regulators, such as insulin pathway components, will have a measurable impact on the ability of *Drosophila* to survive microbial infection. To test this hypothesis, we determined the impact of common fly pathogens on the survival of wild-type or insulin pathway mutant flies. We found that mutations in the insulin pathway significantly impacted host survival and bacterial loads in a manner that depended on the route of infection, and the identity of the infectious microbe. Our results support a model where integrated immune-metabolic activity is critical for host responses to microbial infection.

## RESULTS

### Activation of IMD in the fat body modifies metabolism

Chronic inflammation is a hallmark of metabolic disorders such as type two diabetes. However, we do not fully understand the extent to which host immunity controls metabolic homeostasis. To directly examine the effects of persistent immune signaling on a metabolic organ, we used the R4GAL4 driver line to express a constitutively active IMD (ImdCA) construct exclusively in the fat body (*R4/imdCA*). This approach allowed us to ask how persistent immune activity influences host physiology without collateral damage through the introduction of pathogenic microbes.

Initially, we used whole-genome microarrays to compare the transcription profiles of *R4/imdCA* larvae to control, age-matched *R4GAL4/+* (*R4/+*) larvae. Activation of IMD deregulated the expression of 1188 genes in third instar larvae by a factor of 1.5 or more (Supplemental Tables I and II). We confirmed deregulated expression for six representative genes in subsequent qPCR assays (Fig. 1D). As expected, many response genes, such as antimicrobial peptides, have established roles in the elimination of microbial invaders (Fig. 1A,1B). However, we also noted substantial effects of IMD activation on the expression of genes that control metabolism (Fig. 1A-C, Supplemental Tables I and II). Of the 807 IMD response genes with annotated biological functions, 247 are classified as regulators of metabolic processes (Supplemental Tables I and II). For example, activation of IMD diminished expression of TOR pathway genes (Fig. 1A); decreased expression of *dilp3* (Fig. 1C); increased expression of the insulin pathway antagonists *dilp6*, and *impl2* (Fig. 1C); and elevated expression of the FOXO-responsive transcripts *thor*, and *tobi* (Fig. 1C). Consistent with effects of IMD activation on host metabolism, we observed significant reduction in the expression of enzymes involved in glycolysis, the TCA cycle, mitochondrial ATP production, and fatty acid beta oxidation (Fig. 1A,1B, Supplemental Tables I and II, Supplemental Fig. 1). We also noted that host responses to IMD activation extend beyond the fat body, as IMD activation suppressed the expression of intestinal peptidases and chitin-binding proteins, and lowered the expression of hormone signaling molecules in the salivary glands (Supplemental Tables I). We previously determined the consequences of *imdCA* expression in adult intestinal progenitor cells (24). This allowed us to compare host responses to IMD activation in the fat body, the principle regulator of humoral immunity, to immune activation in the intestine, a first line of defense against oral infection. Interestingly, we observed minimal overlap between the two responses (Supplemental Fig. 2A-1C).

**Figure 1.**
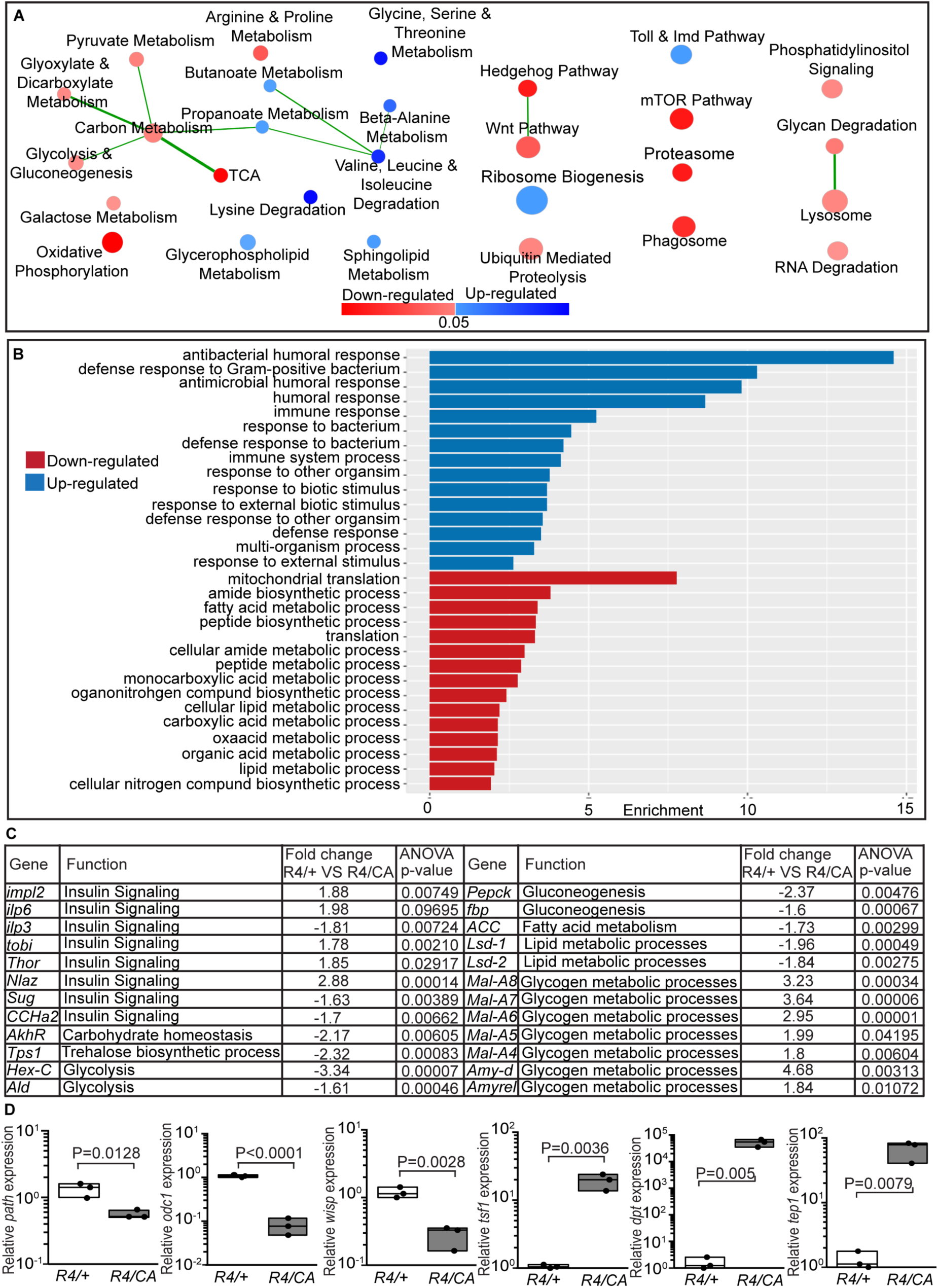
Constitutive IMD activation in the larval fat body alters host biological processes. **(A)** Gene interaction network of upregulated and downregulated KEGG terms altered in *R4/ImdCA* (*R4/CA*) larvae relative to *R4/+* larvae. Red and blue nodes indicate downregulated and upregulated KEGG terms, respectively. Lines indicate genes shared between nodes, and node size indicates the number of genes represented by that KEGG term. **(B)** Biological processes altered in *R4/CA* larvae compared to *R4/+* larvae. Red and blue bars indicate downregulated and upregulated GO terms, respectively. The height of the bar indicates the enrichment score of the GO term. For all terms shown the p value is less that 10^−4^ **(C)** Fold change in the expression of genes involved in insulin signaling, glycolysis, fatty acid metabolism, gluconeogenesis and glycogen metabolism in *R4/CA* larvae relative to *R4/+* larvae. **(D)** Quantification of relative gene expression from third instar *R4/CA* and *R4/+* larvae by qPCR. In each case, gene expression is reported for *R4/CA* flies relative to the corresponding gene in *R4/+* flies. All statistical significance was determined using a Student s t test.

Only 9.8% of genes affected by IMD activation in the fat body were affected by IMD activation in intestinal progenitors (Supplemental Fig. 1A). These observations suggest broad tissue autonomy in IMD responses. In contrast, 29.8% of genes affected by IMD activation are similarly affected by loss of the insulin receptor in fat tissue (25) (Supplemental Fig. 1D, 1E), suggesting an overlap between IMD and insulin-dependent transcriptional outputs in the fat body. This possibility is further supported by a recent examination of the fly transcriptional response to systemic infection with ten distinct bacteria (26). Similar to ImdCA expression, bacterial infection modified the expression of a large number of host genes involved in the regulation of metabolism (Supplemental Fig. 3, Supplemental Table III). Combined, these data sets support a model for the coordinated expression of immune and metabolism regulatory genes in the fat body.

### IMD activation in the fat body disrupts energy reservoirs in the larvae

To test the possibility that IMD activation impacts metabolic homeostasis, we measured carbohydrate levels in *R4/imdCA* and *R4/+* larvae. We did not detect obvious effects of IMD on total glucose levels in larvae (Fig. 2B). However, we found that activation of IMD resulted in significantly higher levels of trehalose, the primary form of circulating carbohydrate in *Drosophila* (Fig. 2A). On average, trehalose levels were approximately twice as high in *R4/imdCA* larvae as *R4/+* larvae (Fig. 2A). We then examined the effects of IMD activation on triglyceride (TG) stores. *Drosophila* larvae store TG in large lipid droplets in the fat body. The lipid storage droplet 1 (Lsd1) and 2 (Lsd2) proteins are involved in lipid storage and lipolysis control, respectively (27), and we found that activation of IMD suppressed expression of both (Fig. 1C). To determine if IMD activation affects TG stores, we measured total TG levels, and lipid droplet size in third instar *R4/imdCA* and *R4/+* larvae. We found that activation of IMD decreased total TG levels by approximately 50% (Fig. 2C), and caused a significant drop in the total volume of lipid stores in the fat body (Fig. 2D, 2E). Together, these results demonstrate that persistent activation of IMD in the fat body causes hyperglycemia, and a depletion of lipid stores.

**Figure 2.**
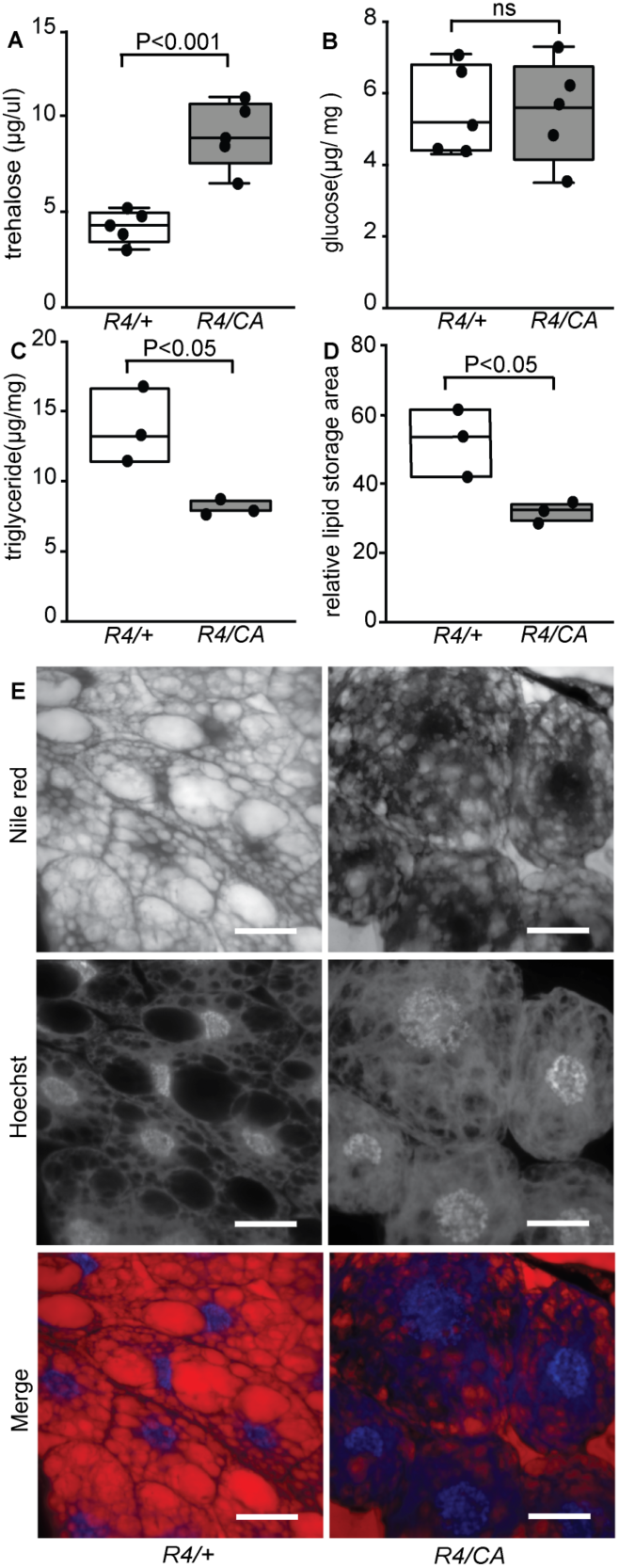
IMD activation disrupts energy reservoirs in the larvae. **(A-C)** Measurement of circulating trehalose (A), total glucose (B) and total triglyceride (C) in *R4/ImdCA* (*R4/CA*) and *R4/+* third instar larvae. **(D)** Quantification of total Nile-red staining area of lipid droplets from third instar larvae after 6 hours starvation. **(E)** Visualization of lipid droplets in third instar *R4/CA* and *R4/+* larvae. Fat tissue was stained with Nile-red (lipid droplets) and Hoechst (nuclei). Scale bars, 25 μm. All Statistical significance tests were performed with a Student’s t test.

### IMD activation in the fat body delays development and reduces pupal size

As nutrient availability influences larval development (28), we tested the effects of IMD activation on larval growth. We were specifically interested in the length of time to pupariation, the size of pupae, and the rate of pupal eclosion, as each of these factors is sensitive to metabolite availability. In each assay, we noted significant effects of IMD activation on development. Specifically, IMD activation delayed the average duration of development from feeding third instar larvae to the P13 stage of pupal development by approximately eighteen hours (Fig. 3A); IMD activation led to a roughly 10% drop in pupal volume (Fig. 3B); and IMD activation caused a significant reduction in adult eclosion rates compared to *R4*/*+* controls (Fig. 3C). In *Drosophila*, dietary nutrients promote cellular and organismal growth by activation of insulin-PI3Kinase-AKT, and TOR-S6 kinase pathways (29). To determine if IMD affects insulin/TOR signaling, we measured the extent of S6 kinase and AKT phosphorylation in *R4/ImdCA* and *R4/+* larvae in four biological replicates. Here, we noticed a significant reduction in the phosphorylation of S6 kinase, and AKT in *R4/ImdCA* larvae relative to *R4/+* controls (Fig. 3D, 3E). Combined, these data indicate that activation of IMD in the fat body disrupts metabolism in *Drosophila*.

**Figure 3.**
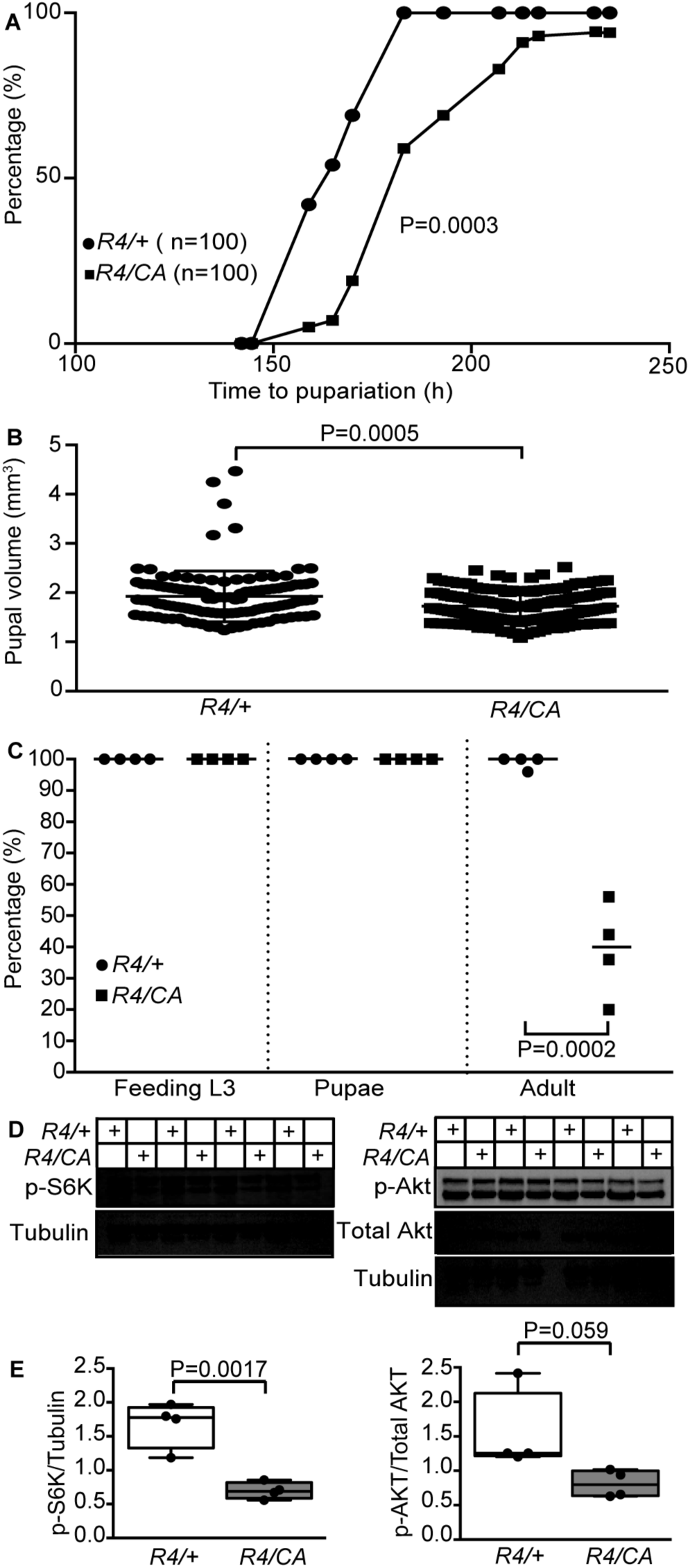
IMD activation impacts larval development. **(A)** Pupariation timing of third instar larvae to P13 stage of pupal development. Statistical significance was determined using a Kolmogorov-Smirnov test, 100 larvae per genotype was used and 4 biological replicates each containing 25 larvae was used in this graph. **(B)** Quantification of pupal volume in *R4/ImdCA* and *R4/+ Drosophila*. Statistical significance was determined using an unpaired student t-test **(C)** 25 feeding third instar larvae of the indicated genotypes were monitored for their development as third instar lavae, pupae, and adults. Results are shown for four independent measurements. **(D)** Western blots of whole lysate from *R4/+* and *R4/CA* third instar larvae probed for p-S6K, p-Akt and total Akt. Tubulin and total AKT levels were visualized as loading controls. **(E)** Quantification of immunoblots of whole lysate from third instar larvae. Statistical tests were performed using an unpaired student t-test.

### Loss of IMD Disrupts Metabolic Homeostasis in *Drosophila*

To determine if IMD contributes to metabolic homeostasis in the absence of an activating signal, we measured insulin expression, body weight, glucose and TG levels in *imd* mutant adult flies raised for ten days on a sugar/yeast (SY) mix, and a high-sugar/yeast version with elevated sucrose levels (SYS). In these assays, we noticed sex-dependent effects of IMD on host-responses to the respective foods. In males, we observed significantly less expression of *dilp2, dilp3*, and *dilp5* in *imd* mutants relative to wild-type controls raised on the SY diet, and significantly less *dilp3* expression in *imd* males raised on the SYS diet (Fig. 4A-C). Furthermore, whereas wild-type males had lower total body weight (Fig. 4E), and higher TG on SYS food relative to wild-type males raised on SY food (Fig. 4G), we did not see any effects of food change on the weight or TG levels of *imd* mutants. Instead, we observed a significant increase in glucose levels in *imd* mutants relative to wild-type controls raised on SYS (Fig. 4F). In contrast to wild-type controls, *imd* mutant females displayed a significant drop in *dilp5* expression levels in flies raised on SYS food relative to flies raised on SY food (Fig. 4J). Likewise, we noticed elevated body weight at day 0 (Fig. 4K), and a significant drop in glucose levels of mutants raised on the SY diet in comparison to the wild-type controls (Fig. 4M).

**Figure 4.**
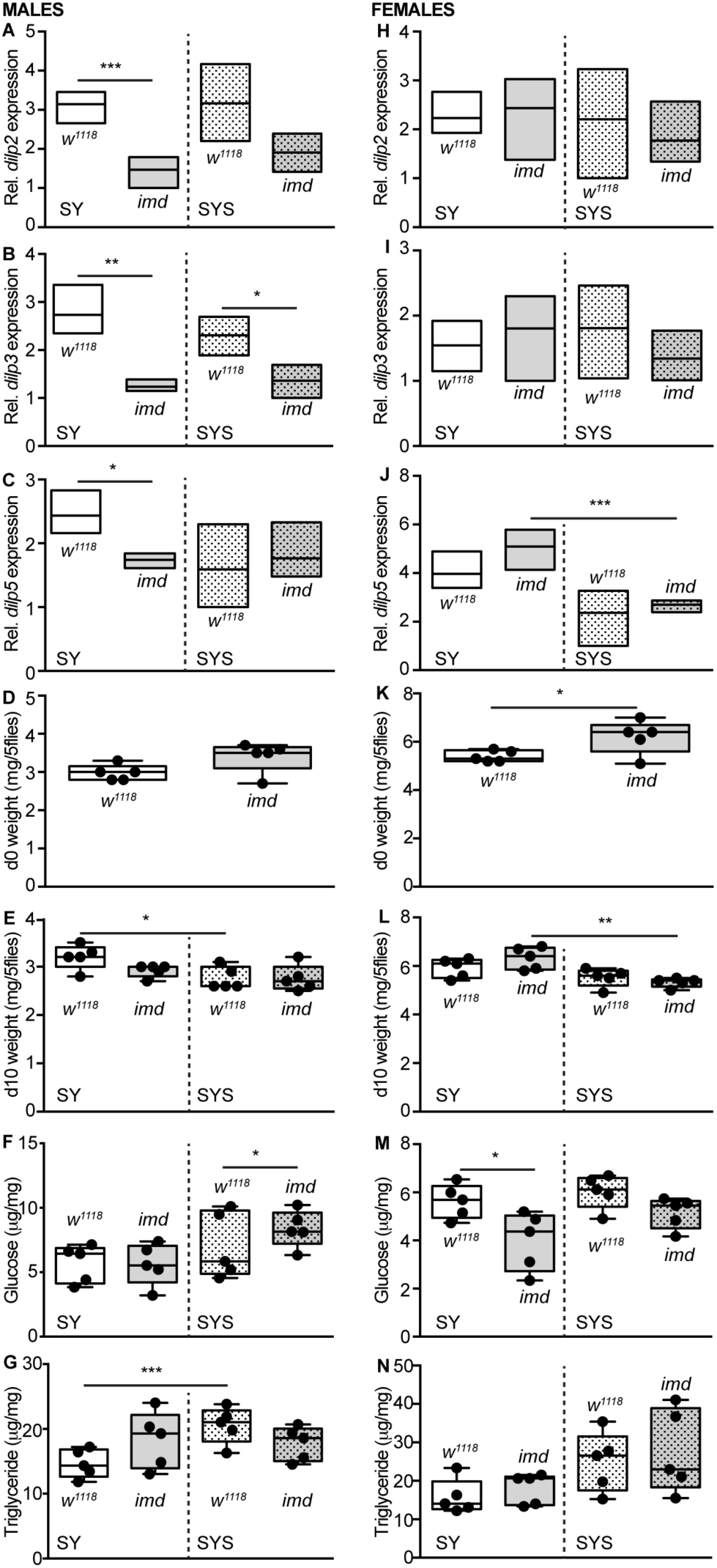
Disruption of the IMD signaling alters metabolic homeostasis. **(A-C, and H-J)**. Quantification of the relative expression of *dilp2* (A, H), *dilp3* (B, I) and *dilp5* (C, J) in *w^1118^* and *imd* mutant male (A-C) and female (H-J) flies raised on sugar/yeast (SY) mix and a high-sugar/yeast version with elevated glucose levels (SYS) for ten days. **(D and K)** Weight measurements of 0–1 day old male (D) and female (K) *w^1118^* and *imd* mutant flies. **(E and L)** weight measurements of male (E) and female (F) *w^1118^* and *imd* mutant flies raised on sugar/yeast (SY) mix and a high-sugar/yeast version with elevated glucose levels (SYS) for ten days. **(F-G and M-N)** Glucose and Triglyceride measurements of male (F-G) and female (M-N) *w^1118^* and *imd* mutant flies raised on sugar/yeast (SY) mix and a high-sugar/yeast version with elevated glucose levels (SYS) for ten days. For all tests, statistical significance were determined using one way ANOVA with Sidak’s correction for multiple comparisons.

To follow the effects of IMD on metabolism more closely, we monitored metabolic activity in adult males raised on a previously described holidic food (30) for twenty days. This food provides all nutrients needed to sustain adult life, and allows investigators to monitor host physiology on a chemically-defined food. We found that mutation of *imd* significantly impairs the expression of *dilp3* (Fig. 5B), diminishes total dILP2 levels (Fig. 5D), and increases the amount of circulating dILP2 (Fig. 5E). As insulin is an essential regulator of glucose metabolism, we determined the effect of *imd* mutation on the ability of one or twenty day old flies to tolerate a glucose meal after a period of fasting. In both cases, we detected significantly higher levels of glucose in mutants immediately upon conclusion of the fast, and during the first two hours after feeding (Fig. 5F,5G). Furthermore, we found that *imd* mutants weighed more (Fig. 5H), and had elevated glucose (Fig. 5I) and TG levels (Fig. 5J) relative to wild-type controls. As *imd* mutants weigh more, we asked if IMD influenced feeding in adult males. To address this question, we raised adult males on holidic food for twenty days, and then quantified their consumption of solid holidic food in a FlyPad assay, or liquid holidic food in a CAFÉ assay. Although we observed an increase in total feeding bouts on solid food (Fig. 5K), mutation of *imd* did not have a significant effect on the total number of sips (Fig. 5l), or on the consumption of liquid food (Fig. 5N), suggesting that IMD does not influence food consumption. In short, our data uncovers a wide range of effects of IMD on insulin, metabolism, and energy storage in adult flies raised on three distinct foods, supporting a role for IMD in the maintenance of metabolic homeostasis.

**Figure 5.**
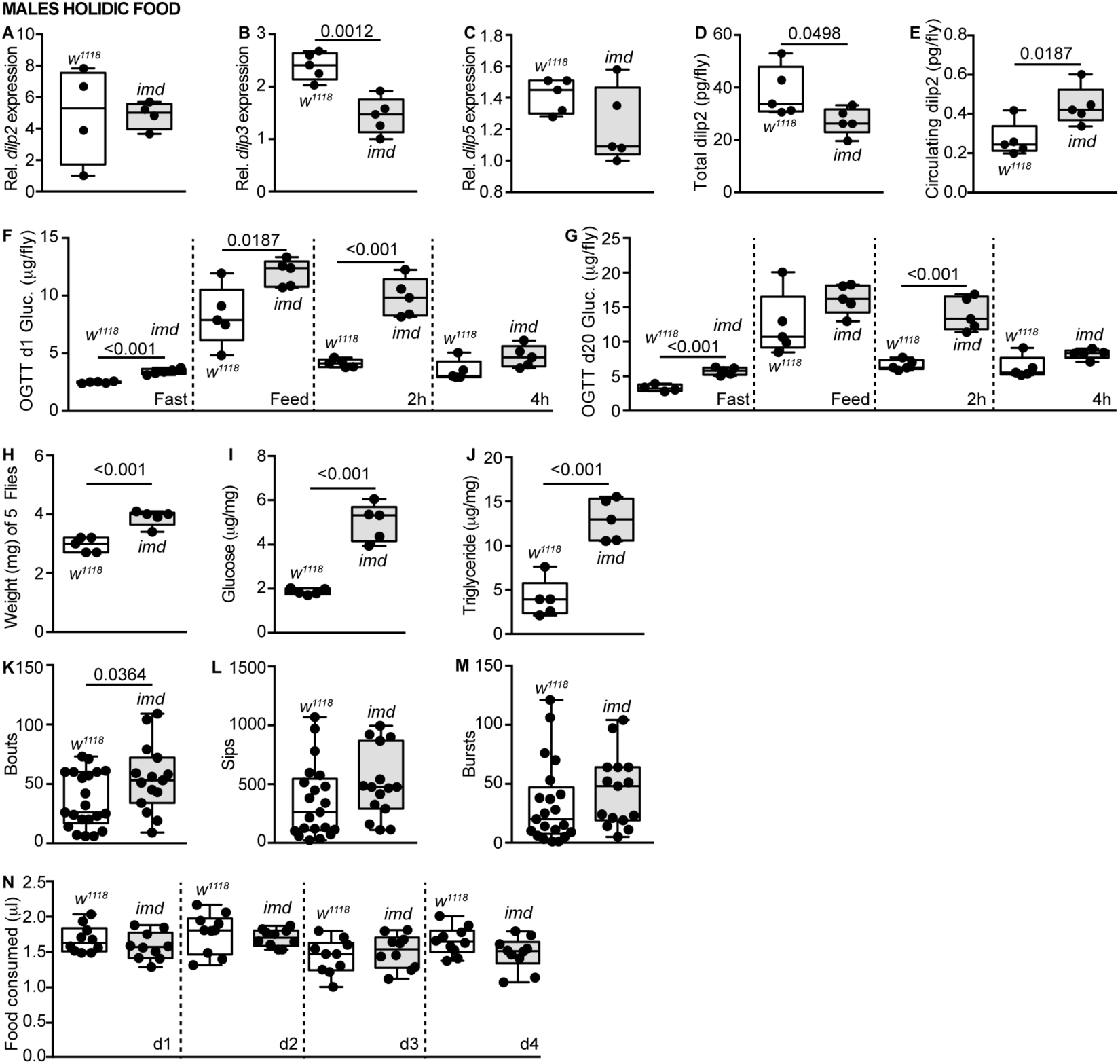
Loss of IMD disrupts metabolism. **(A-E)** Quantification of the relative expression of *dilp2* (A), *dilp3* (B), *dilp5* (C), total DILP2 protein (D) and circulating DILP2 protein (E) in male *w^1118^* and *imd* mutant flies raised on holidic diet. **(F-G)** Oral glucose tolerance test performed on one (F) or twenty (G) day old male *w^1118^* and *imd* mutant flies raised on a holidic diet. **(H-J)** Measurements of weight (H), Glucose (I) and Triglyceride (J) of male *w^1118^* and *imd* mutant flies raised on the holidic diet for twenty days. **(K-N)** Quantification of total feeding bouts (K), number of sips (L) and duration of feeding bursts (M) in twenty days old male *w^1118^* and *imd* mutant flies raised on holidic diet using a FlyPAD. Quantification of liquid holidic food consumption for twenty days old male *w^1118^* and *imd* mutant flies raised on holidic diet using a CAFÉ assay (N). Comparisons were performed using student’s t-tests and p values below 0.05 are indicated throughout.

### Insulin Pathway Mutants Have Bacteria-Specific Responses to Infection

We showed that IMD impacts metabolic homeostasis in both larvae and adult flies. However, it is not clear if metabolic deregulation influences the ability of *Drosophila* to combat bacterial pathogens. As insulin is one of the principle regulators of metabolic homeostasis, we examined the immune responses of *ilp2,3,5* mutant flies to oral and septic challenges with a panel of bacteria that range from low to high pathogenicity in *Drosophila* infection models. For septic infections, we pricked flies in the thorax with a fine needle that deposits the bacteria into the body cavity. As a control, we monitored the survival of flies that we pricked with an uncontaminated, sterile needle. We measured survival and bacterial load as indicators of pathogenicity in our study. In each assay, we found that a sterile wound had minimal effects on the viability of *ilp2,3,5* mutants, or of wild-type flies (Fig. 6A, 6D, 6G, 6K). In contrast, we found that *ilp2,3,5* mutants showed different response towards each pathogen. Systemic infection with *Providencia sneebia*, a microbe that fails to activate IMD (31), did not result in a significant difference between survival of *ilp2,3,5* mutants and control *w^1118^* flies (Fig. 6A). In contrast, the survival of *ilp2,3,5* mutants infected with *Providencia rettgeri, Serratia marcescens Db11*, or *Enterococcus faecalis* was significantly reduced compared to the survival of control *w^1118^* flies (Fig. 6D, 6G, 6K). As *ilp2,3,5* mutants weigh significantly less than wild-type flies (Supplemental Fig. 4A), we determined the bacterial load per fly, and per fly normalized to weight, in the respective infection assays. There was no significant difference between bacterial load in *ilp2,3,5* mutants and *w^1118^* flies six or twelve hours post infection with *P. sneebia* before or after we corrected bacterial load for host weight (Fig. 6B, 6C). In contrast, the bacterial load at 12 hours of septic infection with *P. rettgeri* and *E. faecalis* were significantly higher for *ilp2,3,5* mutants (Fig. 6E,6H). When we corrected bacterial load for host weight, the difference between *ilp2,3,5* mutants and *w^1118^* flies remained unchanged (Fig. 6F,6I). Thus, loss of insulin has microbe-dependent effects on bacterial pathogenicity in *Drosophila*.

**Figure 6.**
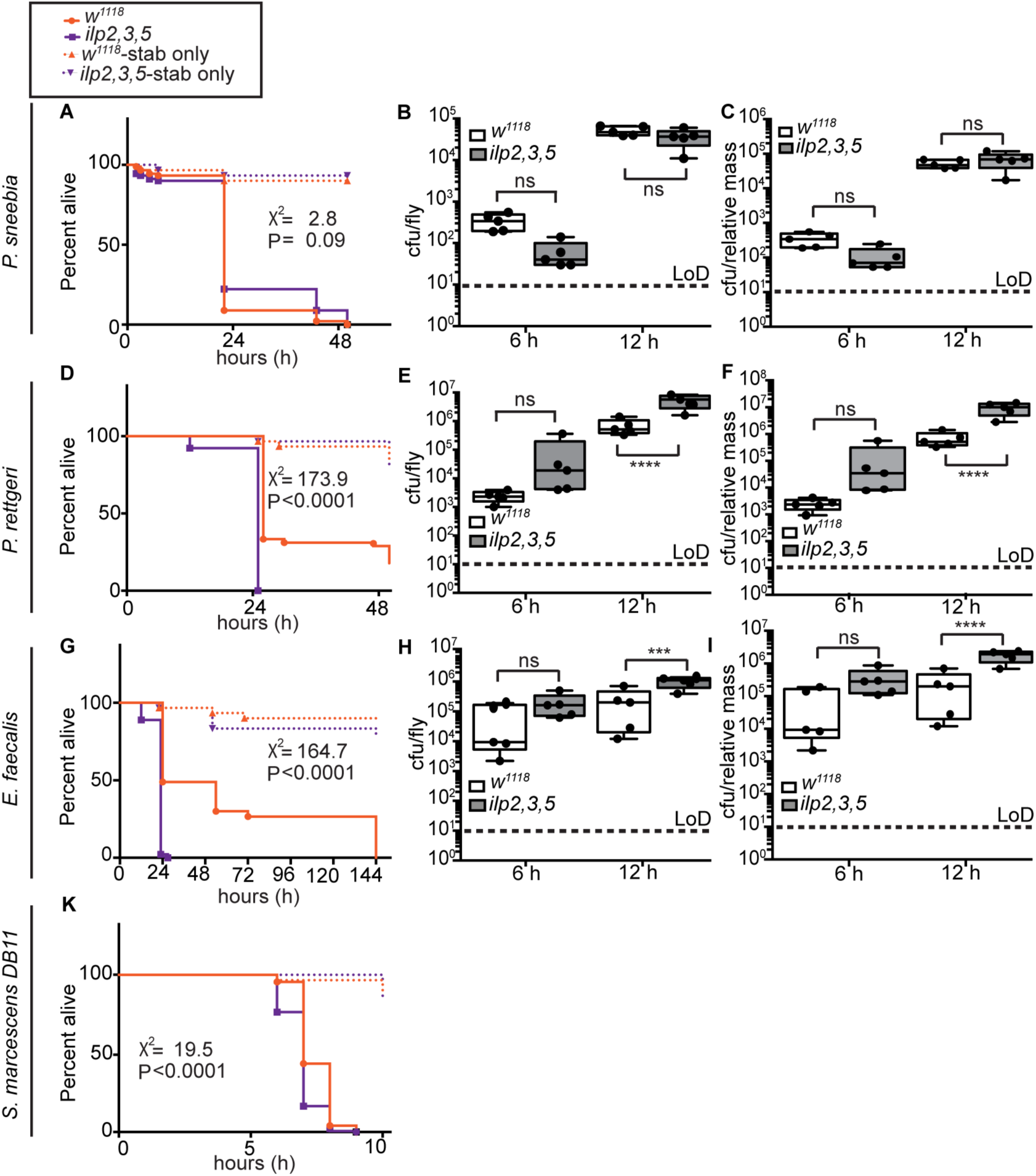
Insulin mutant flies induce a unique response upon septic infection with bacterial pathogens. For all survival curves, solid colored lines indicate the survival of flies of the indicated genotypes challenged with the respective bacteria, and dashed lines indicated the survival of flies challenged with a sterile injury. **(A-C)** Survival curves (A), bacterial load (B) and CFU normalized to weight (C) for *w^1118^* and *ilp2,3,5* mutant flies infected with *Providencia sneebia*. (D-F) Survival curves (D), bacterial load (E) and CFU normalized to weight (F) for *w^1118^* and *ilp2,3,5* mutant flies infected with *Providencia rettgeri*. (G-I) Survival curves (G), bacterial load (H) and CFU normalized to weight (I) for *w^1118^* and *ilp2,3,5* mutant flies infected with *Enterococcus faecalis*. (K) Survival curves for *w^1118^* and *ilp2,3,5* mutant flies infected with *Serratia marcescens Db11*. Statistical significance for survival curves was determined using a Log-rank test of the survival data for *w^1118^* and *ilp2,3,5* mutant flies. A One-way ANOVA was used to compare statistical significance for colony forming units and CFU per relative mass between *w^1118^* and *ilp2,3,5* mutant flies and the Sidak correction method was used for multi-comparisons. Asterisk above the data indicates the statistical significance differences (*= P<0.05, **= P<0.01, ***= P<0.001 and ****= P<0.0001). For all survival experiments, 90 flies per genotypes were used (30 flies in 3 vials) and for CFU measurements, 5 biological replicates each containing 30 flies were used across the infections.

For oral infection experiments, we raised six to seven day-old virgin female flies on cotton plugs soaked with bacterial pathogens, or with LB medium. Bacterial medium alone had minimal effects on fly viability for the first four days (Fig. 7A, 7D, 7G,7J). Here, we found that, with the exception of *E. faecalis* (Fig. 7J), loss of insulin had bacteria-dependent effects on colony forming units (CFU) and host survival. For example, *ilp2,3,5* mutants showed a reduced survival after oral infection with *P. sneebia*, while there was no significant difference between bacterial load in insulin mutant and control flies (Fig. 7A-C). In contrast, we did not observe a significant difference between the survival of *ilp2,3,5* mutants and *w^1118^* flies after oral infection with *P. rettgeri* (Fig. 7D). The bacterial load of *ilp2,3,5* mutants infected with *P. rettgeri* was significantly lower compared to control flies after 48 hours of infection, however, this difference was not significant once we normalized for the weight of *ilp2,3,5* mutants (Fig. 7E, 7F). *ilp2,3,5* mutants lived significant longer than *w^1118^* flies after oral infection with *S. marcescens Db11* (Fig. 7G), and the bacterial load in *ilp2,3,5* mutants was lower than in control flies after 48 hours of infection (Fig. 7H). However, upon correction for weight, we found approximately equal numbers of *S. marcescens Db11* in wild-type and insulin mutant flies (Fig. 7I). Thus, it appears that, controlling for weight, bacterial load is not affected by insulin mutation for each microbe tested. Nonetheless insulin mutation have distinct effects on the ability of the host to survive bacterial challenges. In combination, these observations implicate insulin signaling in the regulation of *Drosophila* immunity to bacterial infection.

**Figure 7.**
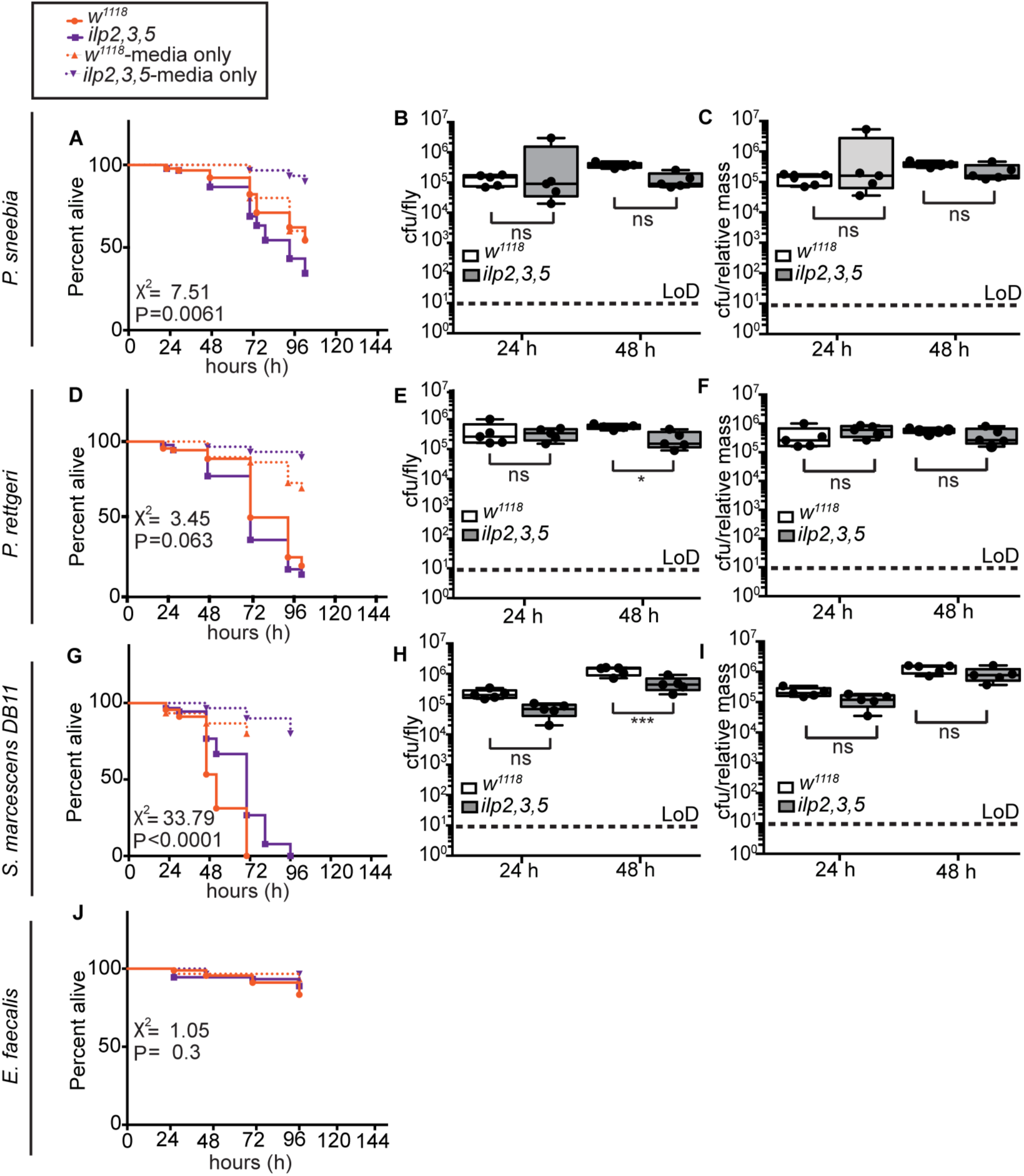
Insulin mutant flies show a diverse response after oral infection with a panel of bacteria. For all survival curves, solid lines indicate the survival of flies of the indicated genotypes challenged with the respective bacteria, and dashed lines indicated the survival of flies control flies fed pathogen-free medium. **(A-C)** Survival curves (A), bacterial load (B) and CFU normalized to weight (C) for *w^1118^* and *ilp2,3,5* mutant flies infected with *P. sneebia*. (D-F) Survival curves (D), bacterial load (E) and CFU normalized to weight (F) for for *w^1118^* and *ilp2,3,5* mutant flies infected with *P. rettgeri*. (G-I) Survival curves (G), bacterial load (H) and CFU normalized to weight (I) for *w^1118^* and *ilp2,3,5* mutant flies infected with *S. marcescens Db11*. (K) Survival curves for *w^1118^* and *ilp2,3,5* mutant flies infected with *E. faecalis*. Statistical significance for survival curves was determined using a Log-rank test of the survival significance between *w^1118^* and *ilp2,3,5* mutant flies. A One-way ANOVA was used to compare statistical significance for colony forming units and CFU per relative mass between *w^1118^* and *ilp2,3,5* mutant flies and then the Sidak correction method was used for multi-comparisons. Asterisk above the data indicates the statistical significance differences (*= P<0.05, **= P<0.01, ***= P<0.001 and ****= P<0.0001). For all survival experiments, 90 flies per genotypes were used (30 flies in 3 vials) and for CFU measurements, 5 biological replicates each containing 30 flies were used across the infections.

### Insulin Modifies Host Immunity to *Vibrio cholerae*

It has been reported previously that enteric infection of *w^1118^* flies with *Vibrio cholerae* suppress insulin signaling (32). To determine if insulin is important for host immunity to *V. cholerae*, we measured host survival and bacterial loads in *Drosophila* that we infected with *V. cholerae*. Here, we found that *ilp2,3,5* mutants have a significantly different survival response towards *V. cholerae* through oral or septic infection. Oral infection with *V. cholerae* resulted in a significantly improved survival of *ilp2,3,5* mutants compared to *w^1118^* flies (Fig. 8A). Improved viability was accompanied by reduced bacterial load in *ilp2,3,5* mutants 48 hours after infection (Fig. 8B). After we normalized the CFUs for host weight, there was still a reduced bacterial load in insulin mutants compared to control flies (Fig. 8C). In contrast, systemic infection with *V. cholerae* lead to a significantly reduced survival in *ilp2,3,5* mutants (Fig. 8G).

**Figure 8.**
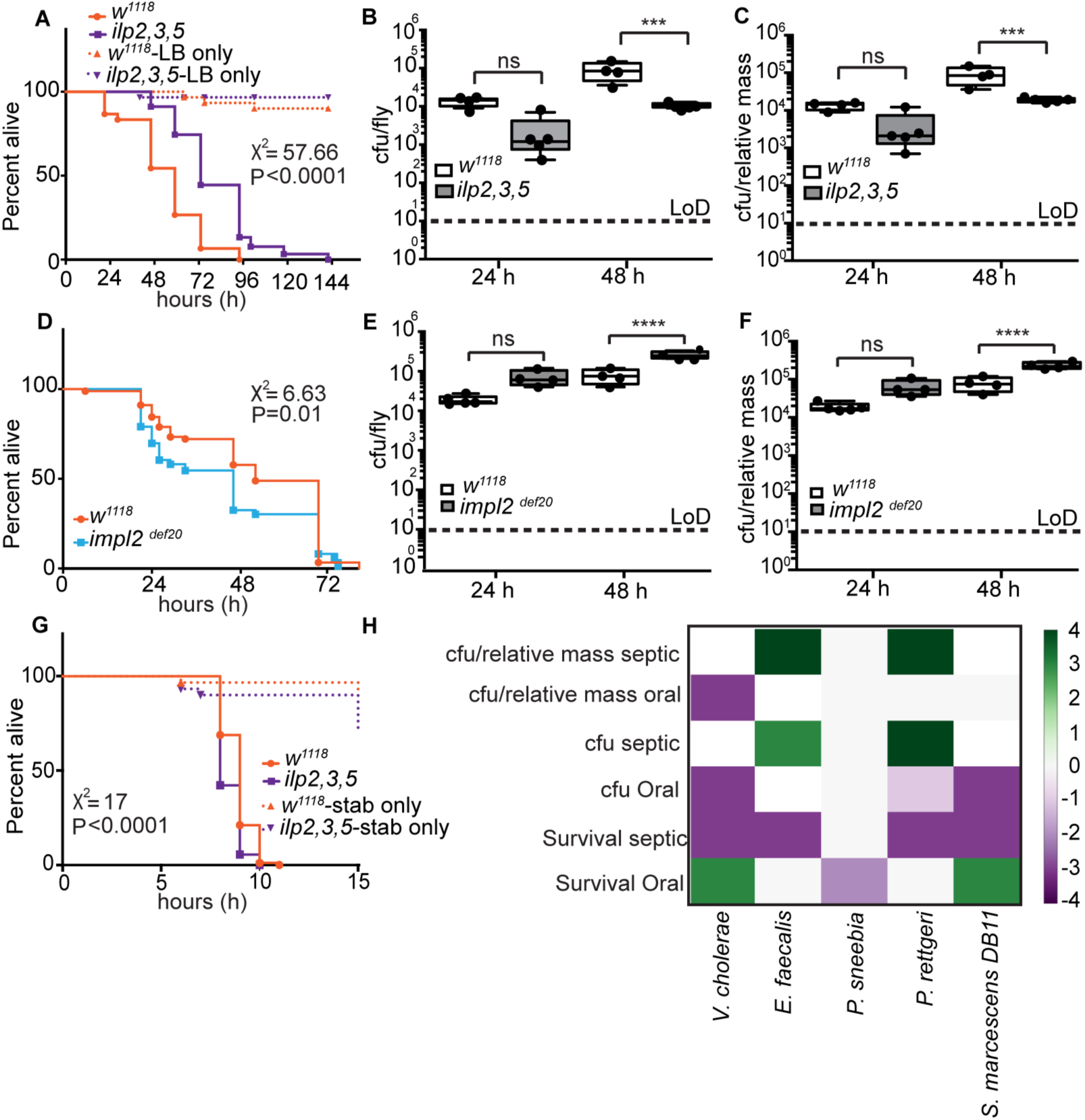
Insulin affects immunity to *Vibrio cholerae*. **(A-C)** Survival curves (A), bacterial load (B) and CFU normalized to weight (C) for *w^1118^* and *ilp2,3,5* mutant flies orally infected with *Vibrio cholerae*. **(D-F)** Survival curves (D), bacterial load (E) and CFU normalized to weight (F) for *w^1118^* and *impl2^def20^* flies orally infected with *Vibrio cholerae*. (G) Survival curves for *w^1118^* and *ilp2,3,5* mutant flies infected with *Vibrio cholerae* through septic infection. Statistical significance for survival curves was determined using a Log-rank test that represent the survival significance between *w^1118^* and *ilp2,3,5* mutant flies and *w^1118^* and *impl2^def20^* flies. One-way ANOVA was used to compare statistical significance for colony forming units and CFU per relative mass between *w^1118^* and *ilp2,3,5* mutant flies and *w^1118^* and *impl2^def20^* flies and then the Sidak correction method was used for multi-comparisons. Asterisk above the data indicates the statistical significance differences (*= P<0.05, **= P<0.01, ***= P<0.001 and ****= P<0.0001). For all survival experiments, 90 flies per genotypes were used (30 flies in 3 vials) and for CFU measurements, 5 biological replicates each containing 30 flies were used across the infections. (H) Heat map summarizing the results of all infections performed in this study. Positive scores indicate experiments where insulin mutants had enhanced survival, or lower bacterial load. Negative scores indicate experiments where insulin mutants had diminished survival, or increased bacterial load. Each experiment was binned according to the degree of significance of the observed phenotype: Scores of 1, or −1 indicate experiments where P<0.05; scores of 2 or −2 indicate experiments where P<0.01; scores of 3 or −3 indicate experiments where P<0.001; and scores of 4 or −4 indicate experiments where P<0.0001.

As insulin deficiency improves host survival after oral challenge with *V. cholerae*, we then asked what effect increased insulin activity will have on the immune response to *V. cholerae*. To answer this question, we examined the immune response of *impl2^def20^* mutants to *V. cholerae*. Impl2 is an antagonist of insulin signaling and *impl2^def20^* mutants contain an amorphic mutation that results in an increased insulin signaling activity. As *impl2^def20^* flies weigh slightly more than *w^1118^* flies (Supplemental Fig. 4B), we also corrected total CFU for weight in these assays. We found that oral infection of *impl2^def20^* flies with *V. cholerae* results in a significantly reduced survival compared to wildtype counterparts (Fig. 8D). The bacterial load in flies with increased insulin signaling was significantly higher compared to control flies (Fig. 8E), and normalization of CFUs for the weight of *impl2^def20^ flies* showed an increased bacterial load in *impl2^def20^* flies compared to *w^1118^* (Fig. 8F). In combination, our data indicate that insulin signaling regulates host immunity to oral infection with *V. cholerae*.

We have summarized the results of all infection assays as a heat map in Fig. 8H. Purples indicate challenges where *ilp2,3,5* mutants have improved survival rates, or diminished bacterial loads, and greens indicate challenges where mutants have an impaired survival, or elevated bacterial loads. These data reveal that loss of insulin affects host survival, and bacterial load in flies challenged with a range of bacterial pathogens. However, the magnitude and nature of the affect depends on the identity of the infectious microbe, and the route of bacterial introduction to the fly.

## DISCUSSION

Eukaryotic life emerged in an environment dominated by microbes and unpredictable availability of nutrients. In response to these challenges, eukaryotes evolved growth and defense responses that support the complexities of multicellular existence. Both responses act in concert to sustain homeostasis, and disruptions to one pathway often impact the other (33). This relationship is particularly evident in insects such as *Drosophila* that rely on their fat body to simultaneously coordinate humoral immunity and nutrient utilization. The fat body detects the nutritional and microbial content of the hemolymph and dictates systemic responses designed to maximize host viability. These dual functions require an integrated response that facilitates the allocation of resources to support growth, reproduction, or antimicrobial responses as needed. The situation is more complex in higher vertebrates where distinct tissues execute metabolic and immune duties. Nonetheless, the organs in question remain in close communication, and metabolic reprogramming is a critical aspect of immune activation in vertebrate lymphocytes (34). Obesity results in recruitment of macrophages to adipose tissue (17, 18), and induces expression of the TNFα adipokine in adipocytes, with consequences for metabolism and inflammation (21). We used *Drosophila* to examine the relationship between IMD activity and metabolic homeostasis. The IMD and TNF signal transduction pathways are closely related, and recent genomic studies implicated IMD in the regulation of metabolism (24, 35, 36). We showed that experimental manipulation of IMD has a substantial effect on host metabolism. Persistent activation of IMD depletes fat reserves, causes hyperglycemia, delays development, and impairs larval growth. In contrast, loss of IMD activity leads to weight gain, elevated storage of TG and glucose, and impaired glucose tolerance. In total, these observations suggest that IMD influences metabolic regulatory pathways. In support of this model, a recent study showed that systemic infection with ten distinct bacterial pathogens has immediate effects on the expression of genes involved in the control of host metabolism (26).

Our findings that IMD affects metabolism corroborates a report that the *Drosophila* TFNα homolog, Eiger, regulates production of insulin peptides in the brain (37), and that the FOXO homolog, Forkhead, regulates intestinal metabolism, and survival after infection in adult *Drosophila* (38). In addition, several studies identified interaction points between immune and insulin responses in the fly. For example, depletion of the insulin receptor from the fat body alters the expression of immune response genes, and alters sensitivity to infection (25). Furthermore, mutations of the IRS homolog *chico* increase survival after infection with *Pseudomonas aeruginosa* and *Enterococcus faecalis* (39); challenges with *Mycobacterium marinum* lower AKT phosphorylation, and diminish systemic insulin activity (14); activation of TOR blocks AMP expression (40); infection increases expression of the FOXO-responsive transcript 4E-BP ortholog *thor* (41); and FOXO regulates the expression of intestinal antimicrobial peptides (42). Combined, these findings suggest a direct relationship between bacterial challenges and insulin-sensitive pathways in the fly. This hypothesis is supported by observations that infection with *M. marinum* lowers triglyceride stores and increases the amount of circulating sugar (14), while challenges with *Listeria monocytogenes* lowers triglyceride and glycogen stores, and inhibits glycolysis (13), and starvation, or protein restriction increases the expression of antimicrobial peptides (43, 44). An earlier study established that activation of the IMD-responsive NF-κB family member Relish fails to affect insulin signaling in the fly (45), suggesting that IMD acts downstream of Relish as a metabolic regulator. We consider the IMD-sensitive c-Jun N-terminal Kinase (JNK) a likely nexus of immune-metabolic control, as several reports link JNK and insulin activity in *Drosophila* (46, 47).

Connections between insulin and immune activity appear conserved through evolution. In *C. elegans*, mutations in the insulin receptor homolog age-1, the PI3-kinase homolog daf-2, or the FOXO homolog daf-16 impact survival after bacterial infection (48, 49). Furthermore, protein restriction improves survival against malaria infections in mice (50), while TFNα regulates glucose and lipid levels in vertebrates (51). Experimentally-induced obesity increases the levels of circulating TFNα (21), and TFNα makes mice less sensitive to insulin signaling, possibly through the regulation of GLUT4 and IRS-1 (52). In humans, nutrient excess leads to an inflammatory state characterized by excess TNF production, and increased insulin resistance (53). Our present work together with these findings indicate that activation of immune responses in a metabolic organ leads to systemic immune-metabolism alterations in the host.

Mechanistically, it is unclear how an immune-metabolic axis influences host responses to bacterial infection. Immunity encompasses resistance mechanisms that kill infectious microbes, and tolerance mechanism that mitigate disease severity without effects on microbial load. In our study, mutations of the insulin pathway affected bacterial load and host survival in a manner that depends on bacterial identity and the route of infection. However, further studies are needed to determine the extent to which insulin regulates tolerance or resistance to the individual pathogens. We consider it likely that the impact of metabolism on host responses to infection is a function of the infectious microbe and the route of infection. For example, glucose supplementation improves survival outcomes in mice challenged with influenza virus, but has the opposite effect on mice infected with *Listeria monocytogenes* (54). This study outlines an accessible model to characterize the relationship between insulin and IMD/TNF-dependent containment of infectious microbes.

## Supporting information

## EXPERIMENTAL PROCEDURES

### *Drosophila* methods

Adult flies and larvae were raised on standard corn meal medium (Nutri-Fly Bloomington Formulation https://bdsc.indiana.edu/information/recipes/bloomfood.html, Genesse Scientific). All adult experiments were performed using virgin male and female flies. For experiments using flies maintained on the holidic diet, the holidic medium was prepared following the published protocol and recipe using the original AA solution (Oaa) at 100mM biologically available nitrogen (Piper *et al*. 2014). The SY diet consists of 0.15M sucrose, 100 g/l (w/v) yeast extract, and 1% (w/v) agar. The SYS diet consists of 1M sucrose, 100 g/l(w/v) yeast extract, and 1% (w/v) agar. Flies that were used in this study are as follows: *w^1118^, R4-GAL4, UAS-ImdCA, impl2^def2^, ilp2,3,5, Ilp2HF* and *imd^EY08573^ (null)* mutants. The *imd* mutants used in this study were back-crossed to the *w^1118^* flies for eight generations prior to use. To measure developmental rates, 25 age-matched feeding third instar larvae were cultured at 25°C and monitored for the formation of wandering third instar larvae, pupae and eclosed adults. For pupariation timing, 25 age-matched third instar larvae were cultured at 25°C and monitored for the length of time required for development to the P13 pupal stage. Developmental and pupariation assays were performed in quadruplicate. For total triglyceride measurement, 10 third instar larvae or 5 adult flies were weighed and homogenized in TE buffer with 01.% Triton X-100. Triglyceride content was measured in larval homogenate using the serum triglyceride determination kit (Sigma TR0100), according to manufacturer’s instructions. Total glucose was measured by homogenizing ten third instar larvae or five adult flies in TE buffer and measuring glucose using the GAGO glucose assay kit (Sigma, GAGO20) according to manufacturer’s instructions. For trehalose hemolymph measurements, groups of 15 third instar larvae were dipped in halocarbon oil 700 (Sigma) and the epidermis was punctured to start hemolymph bleeding. Accumulated hemolymph on the oil drop was aspirated using a glass pipette and immediately frozen on dry ice. 1 μl of hemolymph was mixed with 99 μl trehalase buffer (5mM Tris pH 6.6, 137mM NaCl, 2.7mM KCl) and heated at 70°C for 5 minutes to inactivate endogenous Trehalase. The samples were treated with or without Porcine Kidney Trehalase (T8778–1UN, sigma) and incubated at 37°C for 16 hours, then the reaction was started by adding glucose assay reagent (GAGO20, Sigma), incubated at 37°C for 30 minutes, and the reaction was stopped by adding 12 N sulfuric acid. Absorbance were measured at 540 nm. To calculate trehalose levels, we subtracted glucose levels in untreated samples from glucose levels of samples that were treated with trehalase. CAFE assays were performed as described previously (Diegelmann et al., 2017). We delivered previously mentioned holidic liquid food by leaving out the agar to the capillaries. Each vial contained 3 capillaries with 10 adult flies. Total consumption was calculated every 24 hours for five days. For Nile Red staining, ten third instar larvae were dissected in PBS, and fixed in 4% formaldehyde for 30 minutes. After twice washing with 1X PBS, fat tissues were stained with 1:1000 of a Nile red stock (0.5 mg/ml in acetone) and 1:500 of Hoechst 33258 for 30 minutes. Stained tissue was mounted on slides and visualized using a spinning disk confocal microscope (Quorum WaveFX). Lipid area was quantified with Columbus software (Perkin Elmer). Pupal volume was calculated as previously described (Delanoue et al., 2010). In brief, 24 hr AEL larvae were collected and put into food vials in groups of 50 larvae. Using a paintbrush, 1 day old pupae were picked off the side of the vial. Pupae were imaged using a Zeiss Stereo Discovery V8 microscope using a 14X magnification. Axiovision software was used to measure the length and width of each pupae. Pupal volume was calculated with the assumption that the pupae are cylindrical using the formula: (4/3π)x(length/2)x(diameter/2)^2^.

### FlyPad

We acquired the FlyPad instrument from Dr. Pavel M. Itskov (Itskov *et al*. 2014). We raised male *w^1118^* and *imd* flies on a holidic diet for 20 days. For the FlyPad experiment, we used a holidic medium with agarose substituted for the agar. Prepared food was melted at 95°C and then maintained at 60°C to facilitate pouring. Individual flies were placed in each FlyPad arena with a mouth aspirator at an n=32 for each genotype. Eating behaviour was recorded for 1 hour.

### Oral Glucose Tolerance Test (OGTT)

*w^1118^* and *imd* males were starved overnight for 16 hours on 1% agar, switched to vials containing 10% glucose and 1% agar for 2 hours, and then re-starved on vials of 1% agar. Samples of 5 flies were obtained after initial starvation, after 2 hours on 10% glucose, and then at both 2 hours and 4 hours following re-starvation. Samples of 5 flies were weighed and then mashed in 125 μL TE buffer (10mM Tris, 1mM EDTA, 0.1% Triton X-100, pH 7.4). Glucose was measured using the Glucose Oxidase (GO) Assay kits (Sigma, GAGO20).

### Enzyme-linked Immunosorbent Assay (ELISA)

To measure circulating and total ILP2 levels, we used the *ilp2^1^ gd2HF* fly stock and protocols acquired from Dr. Seung K. Kim (Park *et al*, 2014). For sample preparation, we dissected the black posterior end of the abdomen away and transferred 10 dissected male bodies to 60 μL of PBS, followed by a 10 min vortex at maximum speed. We centrifuged these tubes at 1000Xg for 1 min, and transferred 50 μL of the supernatant to a PCR tube, as our circulating ILP2HF sample. We added 500 μL of PBS with 1% Triton X-100 to the tubes with the remaining flies, and mashed the samples using a pestle and cordless motor (VWR 47747–370), followed by a 5 min vortex at maximum speed. We centrifuged these tubes at maximum speed for 5 min then transferred 50 μL of the supernatant to a PCR tube, as our total ILP2HF sample. For standards, we used FLAG(GS)HA peptide standards (DYKDDDDKGGGGSYPYDVPDYA amide, 2412 daltons: LifeTein LLC). We added 1 μL of the stock peptide standards (0–10 n/ml) to 50 μL of PBS or PBS with 1% Triton X-100. We coated wells of a Nunc Maxisorp plate (Thermo Scientific 44–2404–21) with 100 μL of anti-FLAG antibody diluted in 0.2M sodium carbonate/bicarbonate buffer (pH 9.4) to 2.5 μg/mL, then incubated the plate at 4°C overnight. The plate was washed twice with PBS with 0.2% Tween 20, then blocked with 350 μL of 2% bovine serum albumin in PBS at 4°C overnight. We diluted anti-HA-Peroxidase, High Affinity (clone 3F10) (Roche 12013819001, 25 μg/mL) in PBS with 2% Tween at a 1:500 dilution. We then added 5 μL of the diluted anti-HA-peroxidase to the PCR tubes containing 50 μL of samples or standards, vortexed, and centrifuged briefly. Following blocking, we washed the plate three times with PBS with 0.2% Tween 20. Samples and standards were transferred to wells of the plate, the plate was sealed with adhesive sealer (BIO-RAD, MSB-1001), and then place in a humid chamber at 4°C overnight. Samples were removed with an aspirator and the plate was washed with PBS with 0.2% Tween 20 six times. We added 100 μL 1-Step Ultra TMB – ELISA Substrate (Thermo Scientific 34028) to each well and incubated at room temperature for 30 mins. The reaction was stopped by adding 100 μL 2M sulfuric acid and absorbance was measured at 450 nm on a Spectramax M5 (Molecular Devices).

### Bioinformatics

For microarray studies, we used the GeneChip *Drosophila* Genome 2.0 Array (Affymetrix) to measure gene expression in triplicate assays. Total RNA was extracted from third instar larvae using Trizol. We used 100 ng purified RNA to make labeled cRNA using the GeneChip 3’ IVT Plus Reagent Kit (Affymetrix). We used Transcriptome Analysis Console (TAC) software (Affymetrix) for preliminary analysis of gene expression data. Array data has been submitted to the NCBI GEO database (accession ID: GSE109470). Transcriptome data from *R4/ImdCA* relative to *R4/+* larvae was analyzed using GSEA (55) to identify KEGG pathways that were differentially regulated upon activation of IMD. The data from the GSEA analysis was then visualized using the EnrichmentMap plugin in Cytoscape (version 3.6.1) to generate the gene interaction network (Merico et al. 2010). The resulting network map was curated to remove un-informative nodes, resulting in the simplified network shown in Figure 1A. We used Panther (56) to identify biological process that were affected by IMD activation, and FlyMine (57) to determine tissue enrichment of the respective genes in third instar larvae. The GO term analysis to identify biological processes influenced by *R4/ImdCA* was analyzed using GOrilla (Edan et al. 2009). From the transcriptome data, two gene lists were created that contained significantly upregulated or downregulated genes in response to ImdCA. Each of these lists were run in GOrilla against the background gene set (all microarray genes) with a p-value cutoff of 10^−4^. The top 15 GO terms sorted by p-value were selected for both upregulated and downregulates analyses, ranked by enrichment score, and visualized using the easyggplot2 package in R (version 1.1.442).

### Bacterial Methods

For infection experiments, following bacteria were used: *Providencia sneebia, Providencia rettgeri, Enterococcus faecalis, Serratia marcescens DB 11*, and *Vibrio cholerae* (C6706 strain). For oral infections, all bacteria except *E. faecalis* were streaked from glycerol stocks onto LB plate and grown overnight at 37°C. *E. faecalis* was streaked from glycerol stocks onto brain heart infusion (BHI) plate and grew overnight at 37°C. The following day, we grew single colonies in medium to an OD600 of 0.245, and soaked a sterile cotton plug with 3 ml of the bacterial culture in LB or BHI medium (for *E. faecalis*). 6–7 days old virgin female flies were fed on the cotton plug, and death was recorded at the indicated time points. We used cotton plug soaked with LB medium or BHI medium (for *E. faecalis*) for our control in oral infection experiments. For bacterial load quantification, at indicated time points 25 flies from 5 biological replicates (5 flies from each biological replicates) were collected and surface sterilized by rinsing in 20% bleach, distilled water, 70% EtOH, and distilled water. Then, we randomly distributed these 25 flies into 5 groups (5 flies in each 1.5 ml tube) and then homogenized in respective media. Serial dilutions of fly homogenates were made in 96 well plate and 10 ul of spots were plated on LB agar supplemented with 100 μg/ml streptomycin (to select for *V. cholerae*), BHI agar (to select for *E.faecalis*) and LB-agar for the rest of the bacteria. For calculating CFU per fly, CFU/ml calculated for each bacterial was divided by five. To normalize the CFUs for weight, the CFUS for *ilp2,3,5* flies were divided by the ratio of the average weights of *w^1118^* and *ilp2,3,5* mutants. For septic infection, 0.15mm minutin pins (Fine Science Tools) was dipped into the OD_600_=1 dilution of bacterial which were grown overnight in media at 37°C and then pricked into the thorax of 6-7 days old virgin female flies. A sterile 0.15mm minutin pins was used to prick flies in the thorax and served as control. Flies were then transferred to normal food and kept at 29°C for the rest of the experiments.

### Molecular Techniques

qPCR measurements were performed with RNA purified from whole larvae using Trizol, and the delta delta Ct method was used to calculate relative expression values. For adult flies qPCR measurements, ten heads were homogenized in Trizol. Gene expressions were normalized to *actin*. The following primers were used in this study: *wisp* (F:5‘CAACAACAGTCACTCGTGGG3’, R:5‘TGGAAGAACGAAGATGGTTGC3’), *pathetic* (F:5‘TACTACAGAACTCGCCGCAC3’, R: 5‘CAGACCAAACAGGATGGAGAAC3’), *odc1* (F:5‘ATCTGCGACCTGTCTAGCGT3’, R: 5‘CATTGGATCGTCATTGCACTTG3’), *tep1* (F:5‘AGTCCCATAAAGGCCGACTGA3’, R: 5‘CACCTGCATCAAAGCCATATTG3’), *tsf1* (F:5‘CGATTGTGTGGTGGCTCTGACCAAG3’, R: 5‘AAGGACATCATCCTGAGCCCTCTGC3’), *diptericin* (F:5‘ACCGCAGTACCCACTCAATC3’, R: 5‘ACTTTCCAGCTCGGTTCTGA3’), *dilp2* (F: 5‘TCCACAGTGAAGTTGGCCC3’, R: 5‘AGA TAATCGCGTCGACCAGG3’), *dilp3* (F:5‘AGAGAACTTTGGACCCCGTGAA3’, R:5‘TGAACC GAACTATCACTCAACAGTCT3’), *dilp5* (F:5‘GAGGCACCTTGGGCCTATTC3’, R:5‘CATGTG GTGAGATTCGGAGCTA3’), *actin* (F:5‘TGCCTCATCGCCGACATAA3’, R:5‘CACGTCACCAGGGCGTAAT3’). For Western blots, larvae were lysed in lysis buffer (20 mM Tris-HCl (pH 8.0), 137 mM NaCl, 1 mM EDTA, 25 % glycerol, 1% NP-40, 50 mM NaF, 1 mM PMSF, 1 mM DTT, 5 mM Na_3_VO_4_, Protease Inhibitor cocktail (Roche Cat. No. 04693124001) and Phosphatase inhibitor (Roche Cat. No. 04906845001)), and protein concentrations were measured using the Bio-Rad Dc Protein Assay kit II. For each experiment, equal amounts of protein lysates (usually 15 to 40 μg) were subjected to Western blot analysis. Primary antibodies used were, anti-alpha-tubulin (alpha-tubulin E7, *Drosophila* Studies Hybridoma Bank), anti-phospho-Drosophila Akt Ser505 (Cell Signaling Technology; 4054), and anti-phospho-S6K Thr398 (Cell Signalling Technology; 9209). For immunoblots quantifications, the area under each peak subtracting the background was quantified. The pAkt was normalized to total Akt and the pS6K was normalized to Tubulin.

### Statistical Analyses

All statistical analyses were performed with GraphPad Prism. qPCR data were analyzed with unpaired Student’s t tests (P<0.05). Survival data were analyzed with Log-rank (Mantel-Cox) test. For pupariation timing and pupae counting, Kolmogorov-Smirnov test and unpaired Student’s t tests were used (P<0.05), respectively. Pupal volumes were compared with unpaired Student’s t tests (P<0.05). For analyzing the bacterial load difference, we used one-way ANOVA with Sidak correction.

## AUTHOR CONTRIBUTIONS

S.D., R.D. S.G., and E.F. conceived and designed experiments; S.D., A.G., A.P., R.D. and S.G. performed the experiments; S.D., M.F., R.D., S.G., and E.F. performed data analysis and wrote the paper.

## ACKNOWLEDGEMENTS

*impl2^def20^* was provided by Young Kwon. *ilp2,3,5* and *Ilp2-HF* flies were provided by Seung Kim, respectively. The *imd ^EY08573^* (*null*) mutant flies was provided by Dr. Bruno Lemaitre. The C6706 strain of *Vibrio cholerae* was provided by Dr. Stefan Pukatzki. The following bacterial stocks were provided by Dr. Nicolas Buchon, *E.faecalis, P. sneebia, P. rettgeri* and *S. marcesecens DB11*. FlyPAD was built and assembled by Pavel Itskov. The research was funded by grants from the Canadian Institutes of Health Research to EF (MOP77746) and SG (MOP86622), and from NSERC to SG. MF is supported by an AITF scholarship, and RD by a Clark Smith Brain Tumor Centre Graduate Scholarship. AG was supported by AITF Graduate Scholarship. AP was supported by an AIHS Summer Studentship. We acknowledge microscopy support from Dr. Stephen Ogg at the University of Alberta. Microarrays were processed at the Alberta Transplant Applied Genomics Center.

---

**Figure S1.**
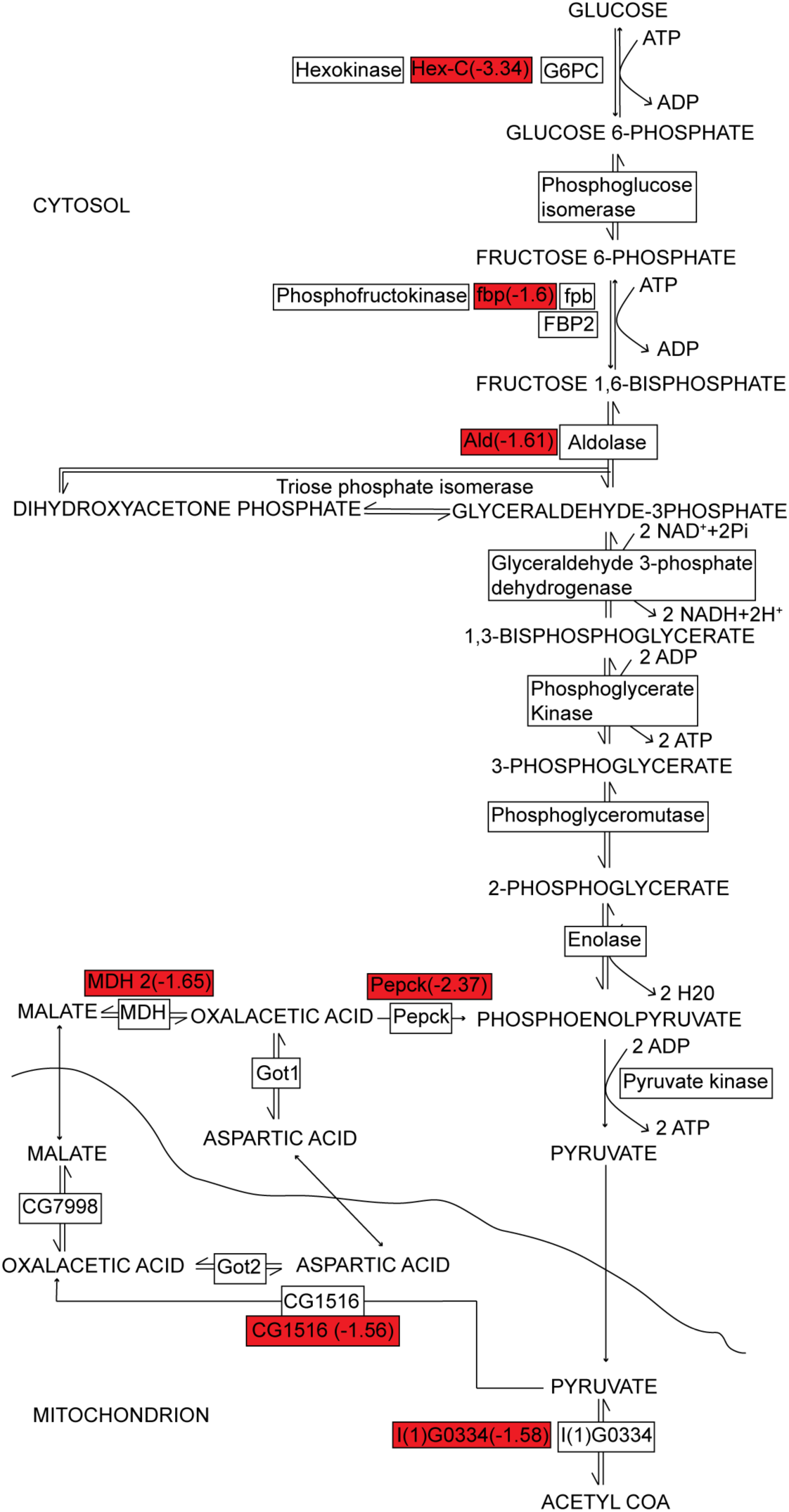
Constitutive IMD activation in the fat body disrupts glycolysis. Red boxes show downregulated enzymes in *R4/ImdCA* larvae relative to *R4/+* larvae, and numbers indicate degree of downregulation.

**Figure S2.**
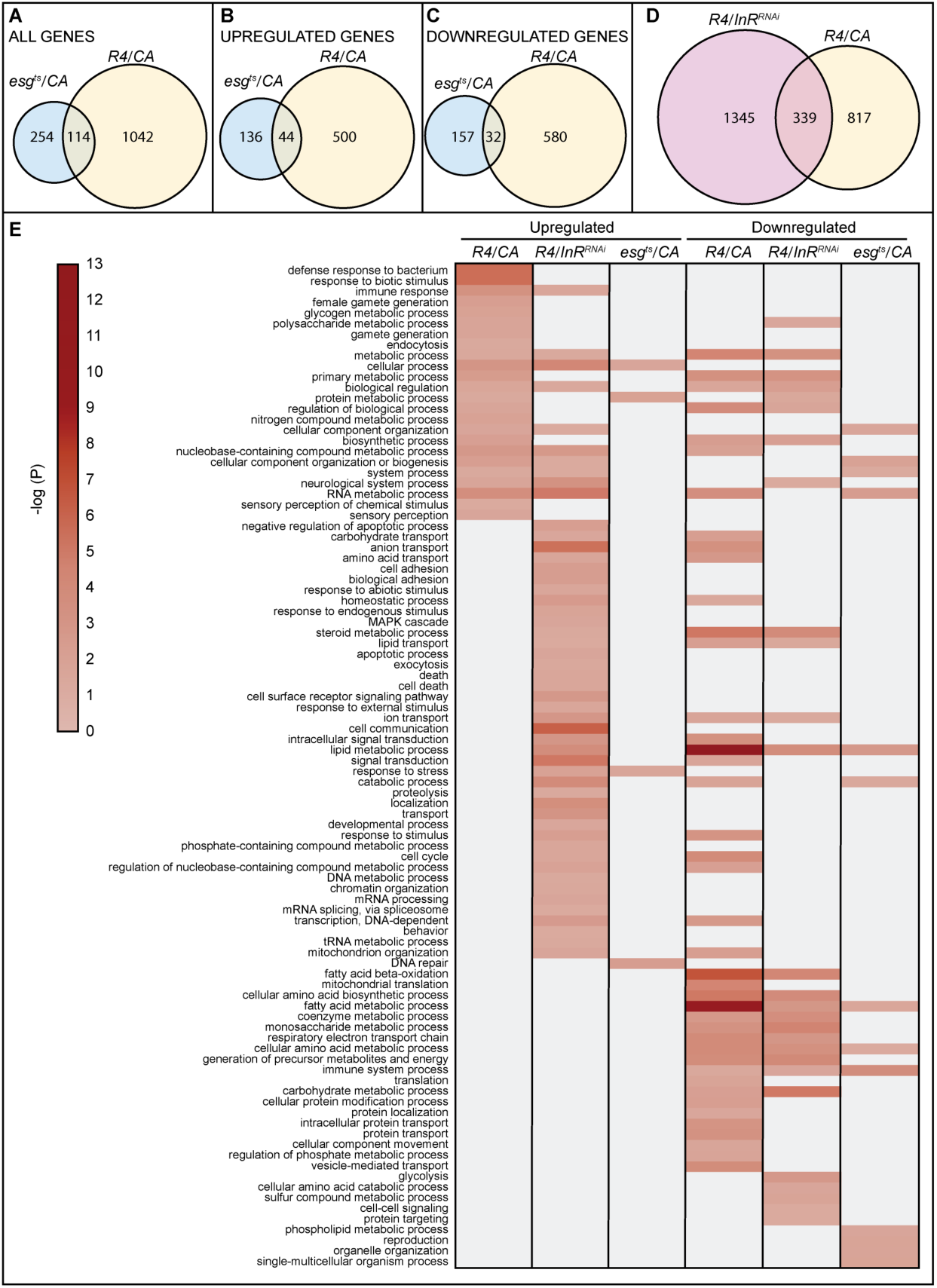
Comparison between constitutive IMD activation in the fat body, intestinal progenitor cells, and larvae with insulin signaling inhibition in the fat body. **(A - C)** Overlap between all dysregulated genes in *R4/ImdCA* larvae and *esg^ts^/ImdCA* intestines. The *esg^ts^* transgenic line allows inducible transgene expression in intestinal stem cells. All genes (A), upregulated genes (B), and downregulated genes (C). **(D)** Overlap between dysregulated genes in *R4/ImdCA* larvae and *R4/InR^RNAl^* larvae. **(E)** Heat-map of dysregulated GO terms in *R4/ImdCA* larvae, *esg^ts^/ImdCA* intestines, and *R4/InR^RNAl^* larvae.

**Figure S3.**
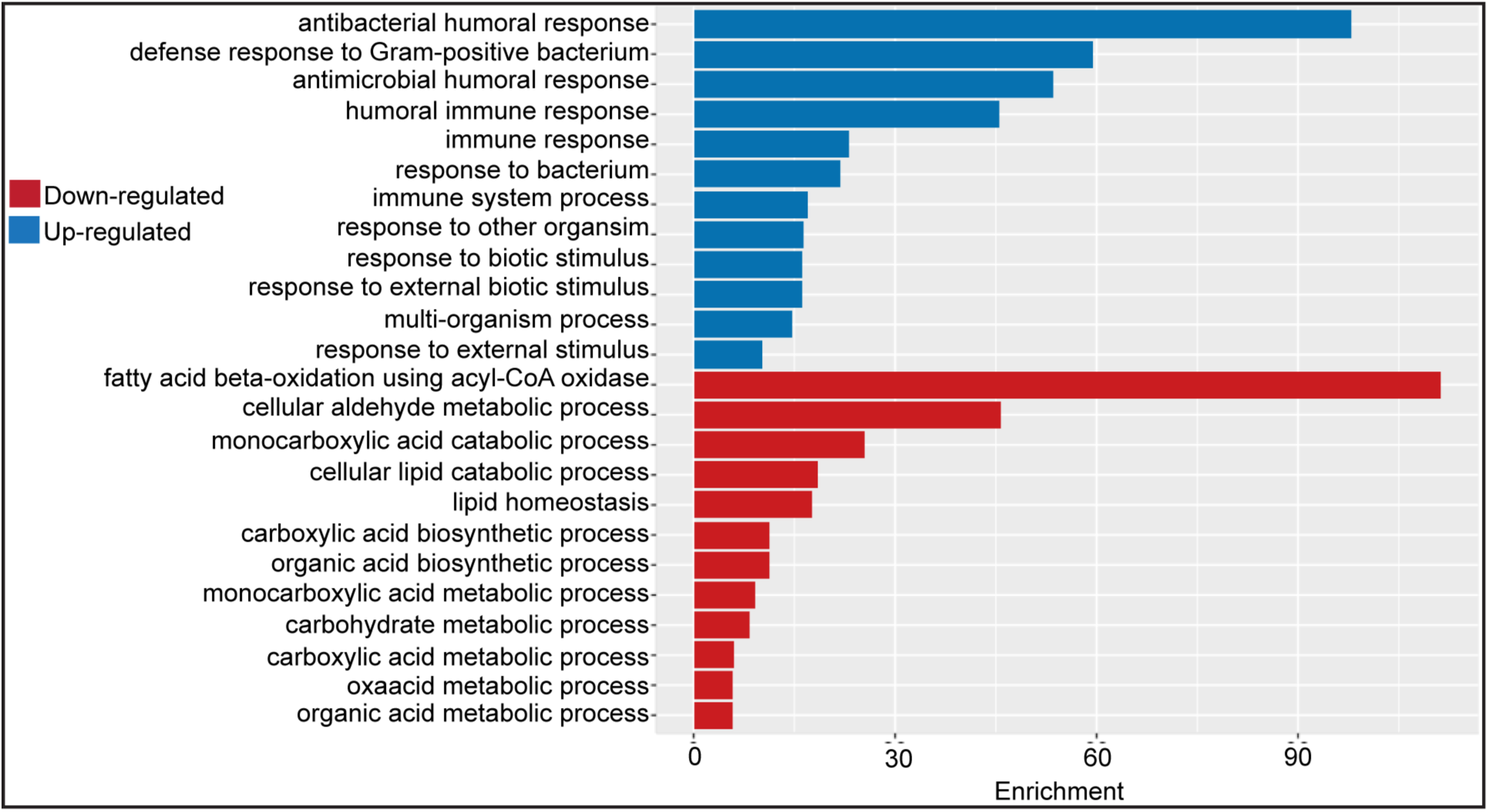
Core biological processes regulated by *R4/CA* and septic bacterial infection. Upregulated and downregulated biological process GO terms from a core list of genes similarly regulated by R4/CA and 7 or more bacterial infections from (26). Red and blue bars indicate downregulated and upregulated GO terms, respectively. The height of the bar indicates the enrichment score of the GO term. For all terms shown the p value is less that 10^−4^.

**Figure S4.**
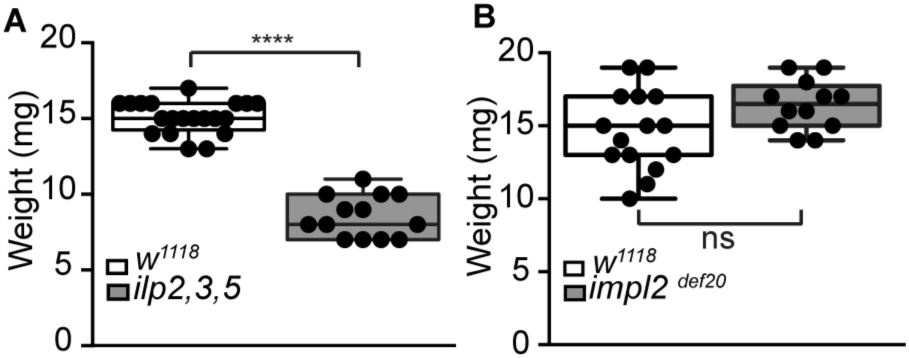
Weight measurements of *w^1118^, ilp2,3,5* and *impl2^def20^* flies. Statistical significance was determined by unpaired student t-test, P< 0.05.

## Supplementary Tables

**Supplementary Table S1.** Description of genes downregulated in *R4/ImdCA* relative *R4/+* larvae are available in R4imdCA_tableS1.

**Supplementary Table S2.** Description of genes upregulated in *R4/ImdCA* relative *R4/+* larvae are available in R4imdCA_tableS2.

**Supplementary Table S3. Comparison between core biological processes regulated by *R4/CA* and septic bacterial infection.** Upregulated and downregulated biological process GO terms from a core list of genes similarly regulated by R4/CA and 7 or more bacterial infections from (26).

